# Motor unit rate coding in intrinsic hand muscles during isolated finger contractions and pinch task

**DOI:** 10.64898/2025.12.19.695509

**Authors:** Nijia Hu, Hélio V. Cabral, Elmira Pourreza, J Greig Inglis, Mikaël Desmons, Francesco Negro

**Affiliations:** Department of Clinical and Experimental Sciences, Università degli Studi di Brescia, Brescia, Italy; School of Physical Education and Sports, Universidade Federal do Rio de Janeiro, Rio de Janeiro, Brazil; Biomedical Engineering Program (COPPE), Universidade Federal do Rio de Janeiro, Rio de Janeiro, Brazil; Postgraduate Program in Rehabilitation Sciences, Universidade Federal do Rio de Janeiro, Rio de Janeiro, Brazil; HAVAE UR 20217, Université de Limoges, Limoges, France

**Keywords:** High-density surface electromyography, first dorsal interosseous, thenar, mean discharge rate, recruitment threshold, motor task

## Abstract

**Purpose:** Muscle force output is modulated via motor unit recruitment and rate coding, yet how rate coding in intrinsic hand muscles differs between isolated and synergistic hand tasks remains unclear. This study examined motor unit discharge behaviour in the first dorsal interosseous (FDI) and thenar during isolated index finger flexion, isolated thumb flexion, and a tip pinch task.

**Methods:** Seventeen participants performed each task at 10%, 20%, and 30% of their maximal voluntary contraction (MVC) while high-density surface electromyograms (HDsEMG) were recorded from both muscles. Motor unit spike trains were completely decomposed from the HDsEMG recordings, tracked across force levels, and their mean discharge rates and recruitment thresholds were calculated.

**Results:** For both FDI and thenar muscles, the mean discharge rate increased with force, but the FDI exhibited steeper slopes than the thenar and at 10-20% MVC than 20%-30% MVC. In addition, lower recruitment thresholds and higher mean discharge rates were observed in the FDI compared to the thenar. Task-dependent differences were also observed in the FDI, with the pinch task yielding higher discharge rates than isolated contractions. In the thenar, differences between tasks were limited to higher forces.

**Conclusion:** These findings demonstrate muscle- and task-specific motor unit modulation across forces in the intrinsic hand muscles, where the FDI relies more on rate coding, while the thenar likely prioritizes recruitment to cope with increased force demands.

## Introduction

In the human neuromuscular system, motor units translate synaptic inputs into muscle contractions, enabling force generation and movement (Heckman & Enoka, 2012). At the motor unit level, force output is modulated through two primary mechanisms: recruitment and rate coding (Enoka & Fuglevand, 2001; Fuglevand et al., 1993). Recruitment involves activating additional motor units, while rate coding refers to adjustments in the discharge rate of already active units (Adrian & Bronk, 1929; Henneman et al., 1965). The relative contribution of these mechanisms during increases in force demand is muscle-dependent (Inglis et al., 2025). For larger muscles like the biceps brachii, recruitment continues throughout most of the force range (∼90% of maximal voluntary contraction, MVC), with rate coding playing a greater role at higher force intensities (Conwit et al., 1999; Kukulka & Clamann, 1981). In contrast, smaller muscles such as the first dorsal interosseous (FDI) typically complete recruitment by 30-50% MVC, after which further increases in force are achieved almost exclusively by discharge rate modulation (Conwit et al., 1999; de Luca et al., 1982). Motor unit discharge behaviour, however, is not determined solely by force level but is also influenced by the motor task being performed, as different motor actions engage different neural inputs and coordination demands (Vaillancourt & Newell, 2003). For example, greater corticospinal excitability, reflected by higher motor evoked potentials, has been observed during precision grasp compared with simple finger flexion (Tinazzi et al., 2003), suggesting that motor units dynamically adjust their discharge behaviour in response to both force and task demands (Avrillon et al., 2024; Mottram, Christou, et al., 2005; Mottram, Jakobi, et al., 2005).

Task-dependent modulation is particularly relevant in the hand, where muscles must generate finely graded forces for dexterous actions of daily living (Ajiboye & Weir, 2009). The intrinsic and extrinsic hand muscles must act in close coordination, such that even apparently isolated finger movements require co-activation of other muscles to stabilize joints and counteract unintended torques (Fuglevand, 2011; Schieber, 1995). Nevertheless, during precise movements such as tip pinch, the intrinsic hand muscles exhibit greater functional independence than the extrinsic muscles, allowing for refined control of fingertip force (Winges et al., 2008). The FDI and thenar muscles are the primary contributors to tip pinch force, as the median nerve (innervating the thenar) block reduces pinch force by ∼60%, whereas the ulnar nerve (innervating the FDI) block produces an ∼85% reduction (Kozin et al., 1999). Biomechanically, their force vectors act in opposite directions but interact synergistically during pinch, with each muscle counterbalancing the other to maintain stability of thumb-index contact (Cooney et al., 1985a; Giurintano et al., 1995). This interplay explains that FDI often shows increased activation during thumb stabilization during isolated contractions and highlights the mechanical coupling that may influence neural control strategies between FDI and the thenar (Hudson et al., 2009). Given that the pinch task is integral to activities such as writing, buttoning a shirt, and lifting delicate objects, clarifying the FDI and thenar motor unit behaviour strategies during these tasks is essential to advancing our understanding of how the central nervous system tailors motor unit control to meet task-specific demands.

Although motor unit behaviour in hand muscles has been widely studied (Adam et al., 1998; Carpentier et al., 2001; Enoka et al., 1989; Fuglevand, 2011; Fuglevand et al., 1999; Kornatz et al., 2005; Spiegel et al., 1996), findings across tasks remain inconsistent, particularly between extrinsic and intrinsic muscle groups (Cabral et al., 2025; Del Vecchio et al., 2023; Tanzarella et al., 2020, 2021). For example, Oßwald et al. (2023, 2025) reported that extrinsic hand flexors recruit different motor units during single-digit versus multi-digit grasping, with higher discharge rates during grasping. In contrast, Maillet et al. (2022) observed no significant task-related differences in discharge rate of intrinsic hand muscles, including the FDI and thenar, between pinch and rotation tasks. Such discrepancies may stem from variations in force levels across studies, which also influence recruitment thresholds and discharge rates of the involved motor units (Carpentier et al., 2001; McNulty et al., 2000). Consequently, disentangling the influence of task demands from force-dependent changes is crucial to understanding neural control in intrinsic hand muscles.

This study aimed to investigate motor unit discharge behaviour of intrinsic hand muscles, specifically the FDI and thenar, during isolated index finger flexion, isolated thumb flexion, and a synergistic pinch task. High-density surface electromyography (HDsEMG) was utilized to record the myoelectric signal, which was subsequently decomposed into motor unit spike trains. These motor unit spike trains were tracked across submaximal force levels (10%, 20%, and 30% MVC) to examine task- and muscle-dependent differences in rate coding strategies. Mean motor unit discharge rate and recruitment thresholds were analysed during the recruitment, plateau, and de-recruitment phases to determine adaptations in neural control following increases in force demands in the FDI and thenar when acting either independently or synergistically. It was hypothesized that motor unit discharge behaviour would differ between the FDI and thenar muscles and could be further modulated by task demands.

## Methods

### Participants

Seventeen healthy adults (6 females, age: 29 ± 5 yr) participated in this study. All participants were right-hand preferred, which was determined by the participants indicating which hand they preferred to write with (Adamo & Taufiq, 2011). No participants reported any history of upper-limb musculoskeletal or neurological disorders. Prior to the experiment, each subject provided written informed consent. The study was approved by the local ethics committee (NP5890) and was conducted in accordance with the ethical guidelines set forth in the latest version of the Declaration of Helsinki.

### Experimental design and protocol

Participants visited the laboratory on one occasion, with the experimental session lasting approximately one and a half hours. Participants were seated comfortably in front of a custom-made manipulandum with their right arm and wrist stabilized on a forearm jig. The elbow was flexed at 80° (0° being anatomical position) and the wrist was positioned in anatomical neutral position (midway between full supination and full pronation). The index finger and thumb were fixed to adjustable supports **(Figure 1A**), which were connected to precision load cells (SM S-TYPE-100N, Interface, U.S. & METRIC) to record the isometric forces produced by flexion of index finger and thumb, either separately or combined during the pinch task.

**Figure 1.**
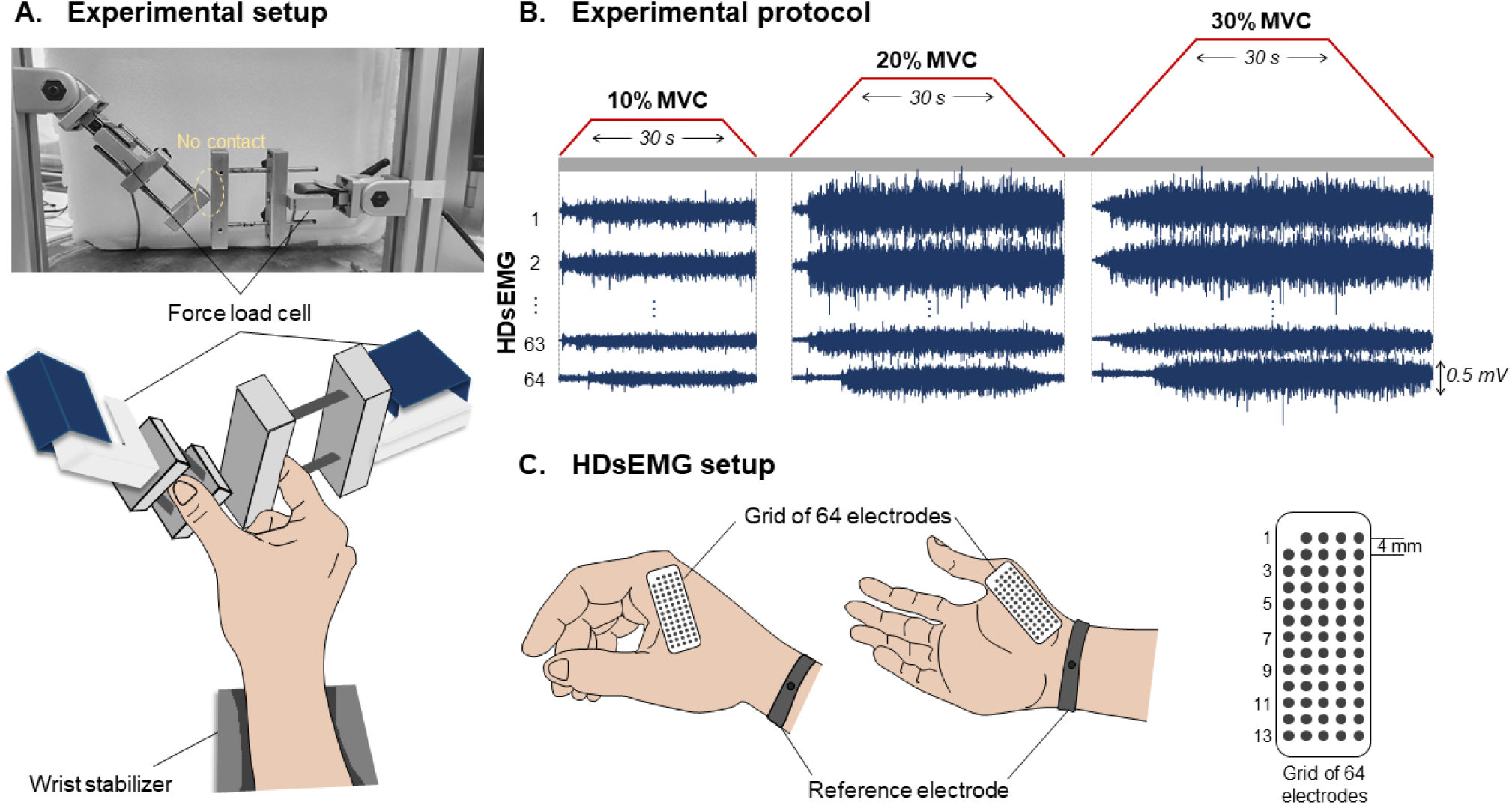
A) The custom-made manipulandum including two load cells to record the isometric forces produced by the flexion of index finger and thumb (upper), and the hand position during experiments (lower). B) The experimental protocol included performing isometric contractions at 10%, 20%, and 30% of the maximal voluntary contraction (MVC) for all tasks (index finger flexion, thumb flexion, pinch). Target trajectories are shown in red, and examples of recorded high-density surface electromyograms (HDsEMG) are displayed in dark blue. C) The position of HDsEMG grids on the first dorsal interosseous (FDI) and thenar muscles (left), and the 64-channel electrode grid arrangement (right).

To determine the maximum isometric voluntary contraction (MVC) of index finger flexion and thumb flexion, participants performed maximal isometric contractions for 5s each. Two trials were performed for each task, and the highest value across trials was used as the reference to calculate the target force levels. After a 2 min rest period, participants were instructed to perform submaximal isometric contractions at 10%, 20%, and 30% MVC following a trapezoidal profile (**Figure 1B**). This profile consisted of three distinct phases: 1) a gradual ramp-up phase from 0% MVC to the target force at a rate of 3% MVC/s; 2) a sustained plateau phase where the force was maintained at the target force for 30 seconds; and (3) a ramp-down phase, which returned from the target force to 0% MVC at a rate of 3% MVC/s (Lulic-Kuryllo et al., 2021; Pollet et al., 2022). Each participant performed two trials for each target force level. The experimental protocol was repeated for each of the three tasks: index finger flexion, thumb flexion, and synergistic flexion of the index finger and thumb (i.e., pinch). The order of the tasks was randomized, with a 1-min rest interval between tasks to prevent fatigue. Real-time visual feedback of the exerted force and trapezoidal target was provided on a monitor positioned at approximately 70 cm in front of the participant.

### Data collection

During all isometric submaximal contractions, HDsEMG signals were recorded from the FDI and thenar using two grids of 64 electrodes each (13 rows × 5 columns, gold coated, 1 mm diameter, 4 mm interelectrode distance, OT Bioelettronica, Turin, Italy; **Figure 1C**). The position and orientation of the grids was determined via palpation by an experienced investigator (EP, MD). Specifically, the grids were attached over the belly of each muscle in the following locations: FDI, lateral to the line connecting the heads of the first and second metacarpals; thenar, on the muscle belly of thenar (Germer et al., 2021) (**Figure 1C**). HDsEMG grids were fixed to the skin using a bi-adhesive foam, and the electrode-skin contact was ensured by filling the foam cavities with conductive paste (AC cream, Spes Medica, Genova, Italy). Reference electrodes were positioned on the right wrist. Before electrode placement, the hand was shaved and skin cleaned with an abrasive paste (EVERI, Spes Medica, Genova, Italy) to reduce electrode-skin impedance and enhance signal quality. HDsEMG signals were recorded in monopolar mode and digitized at 2,000 samples/s using a 16-bit amplifier (10–500 Hz bandwidth; Sessantaquattro+, OT Bioelettronica, Turin, Italy). Force signals measured by the load cell were amplified by a factor of 100 (Forza-j, OT Bioelettronica, Turin, Italy). HDsEMG, force feedback (i.e., trapezoidal profile) and force output were recorded synchronously using the Sessantaquattro+ amplifier.

### Data analysis

#### Motor unit decomposition

From the two performed trials for each submaximal force level, the trial with the lowest root-mean-square error between the force and target signals (i.e., highest force-target matching) was used for further analysis. First, HDsEMG underwent preprocessing, including bandpass filtering with a third-order Butterworth filter (20-500 Hz cutoff frequencies) and visual inspection of monopolar and single-differential signals. During visual inspection, channels with low quality or artifacts were excluded from further analysis. Subsequently, individual motor unit spike trains were decomposed from the HDsEMG signals using a convolutive blind source separation algorithm (Negro et al., 2016). This method has been validated and previously applied to assess the activity of single motor units of the FDI and thenar (Cabral et al., 2024; Negro et al., 2016). Each force trial (10%, 20%, and 30% MVC) was decomposed separately. After the automatic identification of motor units, all the motor unit spike trains were visually inspected for misidentified or missed spikes (Hassan et al., 2020). Spikes leading to non-physiological discharge rates were manually corrected by an experienced investigator through an iterative process, which has been shown to be highly reliable across operators (Hug et al., 2021). Only motor units with a silhouette value above 0.87, which is a metric to evaluate decomposition accuracy (Negro et al., 2016), were included in further analysis.

#### Motor unit tracking

After editing the motor unit spike trains, the motor units were tracked across force levels. To maximize the number of successfully tracked units, this procedure was performed in two steps: first between 10-20% MVC and then between 20-30% MVC. For tracking, the motor unit separation vectors obtained from each individual contraction were reapplied to the concatenated HDsEMG signals of the corresponding pair of contractions (Oliveira & Negro, 2021). For example, separation vectors from motor units decomposed at 10% and 20% MVC were applied to the concatenated signals of 10% and 20% MVC trials, and the same procedure was used for tracking between 20% and 30% MVC trials (**Figure 2A and 2B**). Because separation vectors from both contractions were used on the concatenated signals (**Figure 2C**), motor units recruited at both force levels were inevitably identified twice. Duplicated motor units were identified as having higher than 30% of common discharge times (Holobar et al., 2014), and in such cases, the unit with the higher coefficient of variation of the inter-spike interval was removed from further analysis. To analyse motor units tracked across all three force levels, we identified common motor units at 20% MVC that were matched with both 10% and 30% MVC. Specifically, the motor unit action potential shapes at 20% MVC previously matched with 10% MVC were cross-correlated with those at 20% MVC matched with 30% MVC. Motor units with a high degree of action potential similarity (cross-correlation > 0.8; Martinez-Valdes et al., 2017) were considered to represent the same unit across all three contractions.

**Figure 2.**
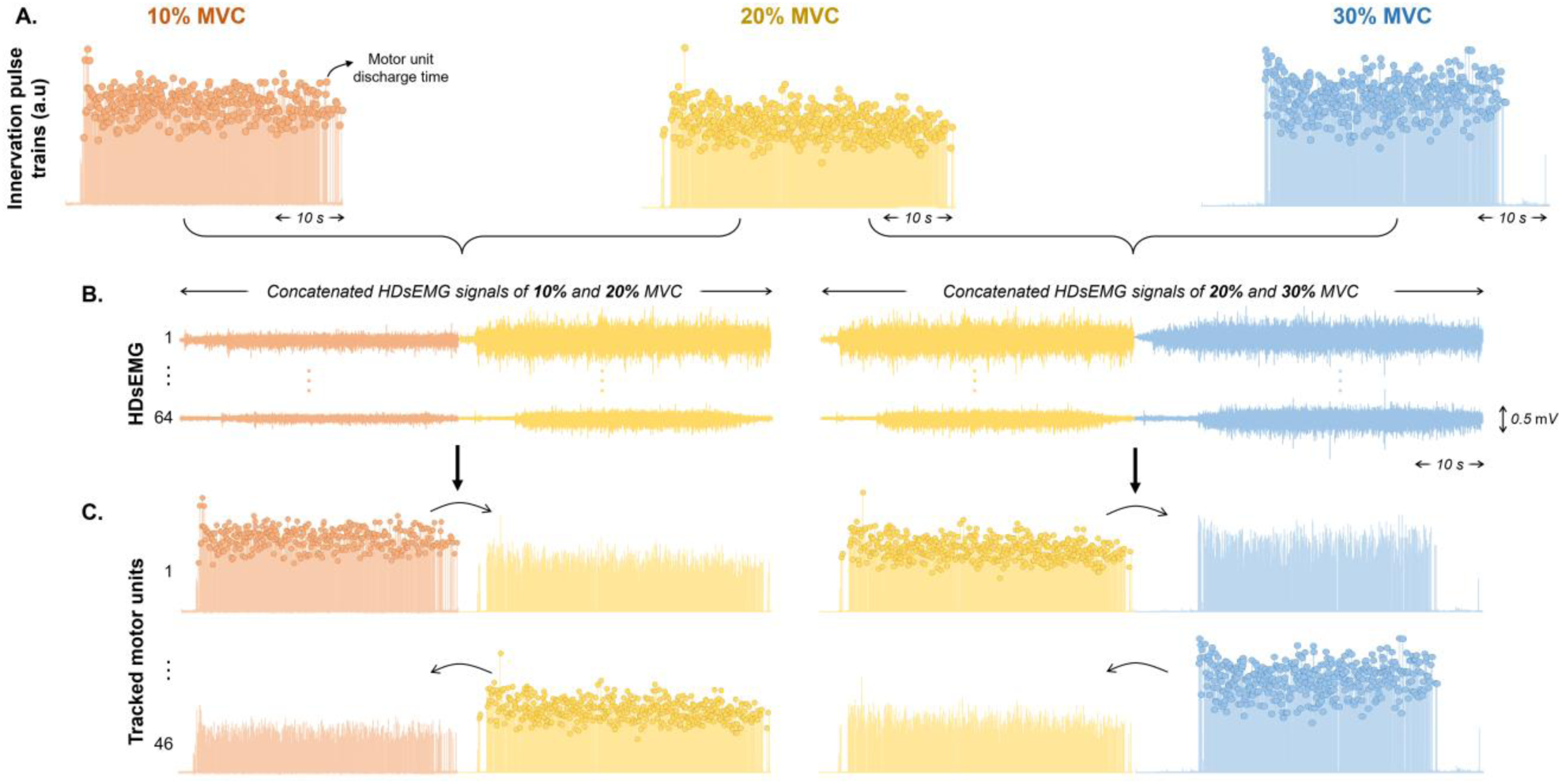
A) HDsEMGs were decomposed separately for each individual force level (10%, 20% and 30% MVC), resulting in innervation pulse trains containing the motor unit discharge times. B) For tracking the motor units across force levels, the motor unit separation vectors obtained from each individual contraction were reapplied to the concatenated HDsEMG signals of the corresponding pair of contractions (10-20% MVC and 20-30% MVC). C) The reapplication of the motor unit separated vectors in the concatenated signals resulted in the tracking of the same motor unit across the corresponding pair of contractions.

#### Motor unit discharge behaviour across force levels

For quality control, after editing, only motor units with silhouette (SIL) > 0.90 (Negro et al., 2016) were considered for further analysis. For each matched motor unit, the mean discharge rate was calculated separately for three phases: recruitment phase (force ascending), plateau phase (steady force), and de-recruitment phase (force descending), respectively. During recruitment and de-recruitment phases, the mean discharge rate was calculated from the first and last two firings. Motor unit recruitment threshold and de-recruitment threshold were calculated as the force value (%MVC) corresponding to the first and last motor unit discharges (Girts et al., 2020).

### Statistical analysis

Statistical analysis was performed using JASP (0.19.3.0). Linear mixed models (LMMs) were used to compare the mean discharge rates and recruitment thresholds across different force levels and tasks. LMMs were also used to compare the mean discharge rate slopes across force between muscles and tasks. For all parameters, random intercept models were applied, with force level (10%, 20%, and 30% MVC), task (isolated and pinch), and muscle (FDI and thenar) as fixed effects, and subject as a random effect. LMMs were implemented using the Kenward–Roger method to approximate degrees of freedom and compute the p-values. In cases of significant interactions, Tukey’s post hoc tests were performed for pairwise comparisons. Estimated marginal means and pairwise comparisons, along with their 95% confidence intervals. Statistical significance was set at α = 0.05. Results are reported as mean ± standard deviation in the text and figures. All individual data of matched motor unit discharge times recorded during each task and force level are available at https://doi.org/10.6084/m9.figshare.30920849.

## Results

### Motor unit yield

**Table 1** provides the average number of motor units decomposed and matched motor units across force levels and tasks. Overall, a higher number of motor units were decomposed and matched in FDI compared to the thenar for both isolated and pinch tasks.

**Table 1.** The average number of decomposed and matched motor units per participant, separately per muscle and task.

|  |  | ISOLATED TASK |  |  | PINCH TASK |  |  |
| --- | --- | --- | --- | --- | --- | --- | --- |
| FDI | Decomposed | 10% MVC | 20% MVC | 30% MVC | 10% MVC | 20% MVC | 30% MVC |
|  |  | 13 ± 7 | 15 ± 8 | 15 ± 7 | 15 ± 7 | 15 ± 9 | 17 ± 8 |
|  | Matched | 10-20% | 20-30% | 10-20-30% | 10-20% | 20-30% | 10-20-30% |
|  |  | 9 ± 5 | 11 ± 5 | 5 ± 3 | 6 ± 3 | 10 ± 5 | 11 ± 6 |
| THENAR | Decomposed | 10% MVC | 20% MVC | 30% MVC | 10% MVC | 20% MVC | 30% MVC |
|  |  | 7 ± 4 | 8 ± 6 | 10 ± 6 | 8 ± 7 | 9 ± 7 | 10 ± 7 |
|  | Matched | 10-20% | 20-30% | 10-20-30% | 10-20% | 20-30% | 10-20-30% |
|  |  | 4 ± 2 | 5 ± 4 | 2 ± 1 | 3 ± 3 | 4 ± 4 | 2 ± 2 |

### Motor unit mean discharge rate

#### Matched units between 10-20% and 20-30% MVC

During the plateau phase, mean discharge rate significantly increased with force in the FDI, from both 10% to 20% MVC (F_(1,608.01)_ = 403.006, p < 0.001), and 20% to 30% MVC (F_(1,700.01)_ = 219.862, p < 0.001, **Figure 3A**). Similar results were observed in the thenar, with significant increases from 10% to 20% MVC (F_(1,191.49)_ = 14.283, p < 0.001) and 20% to 30% MVC (F_(1,286.27)_ = 29.478, p < 0.001, **Figure 3B**). In the FDI, the increase in mean discharge rate was significantly steeper from 10% to 20% MVC compared to 20% to 30% MVC (p < 0.001, **Figure 3C**), whereas no differences in slope were observed in the thenar (p = 0.132, **Figure 3D**).

**Figure 3.**
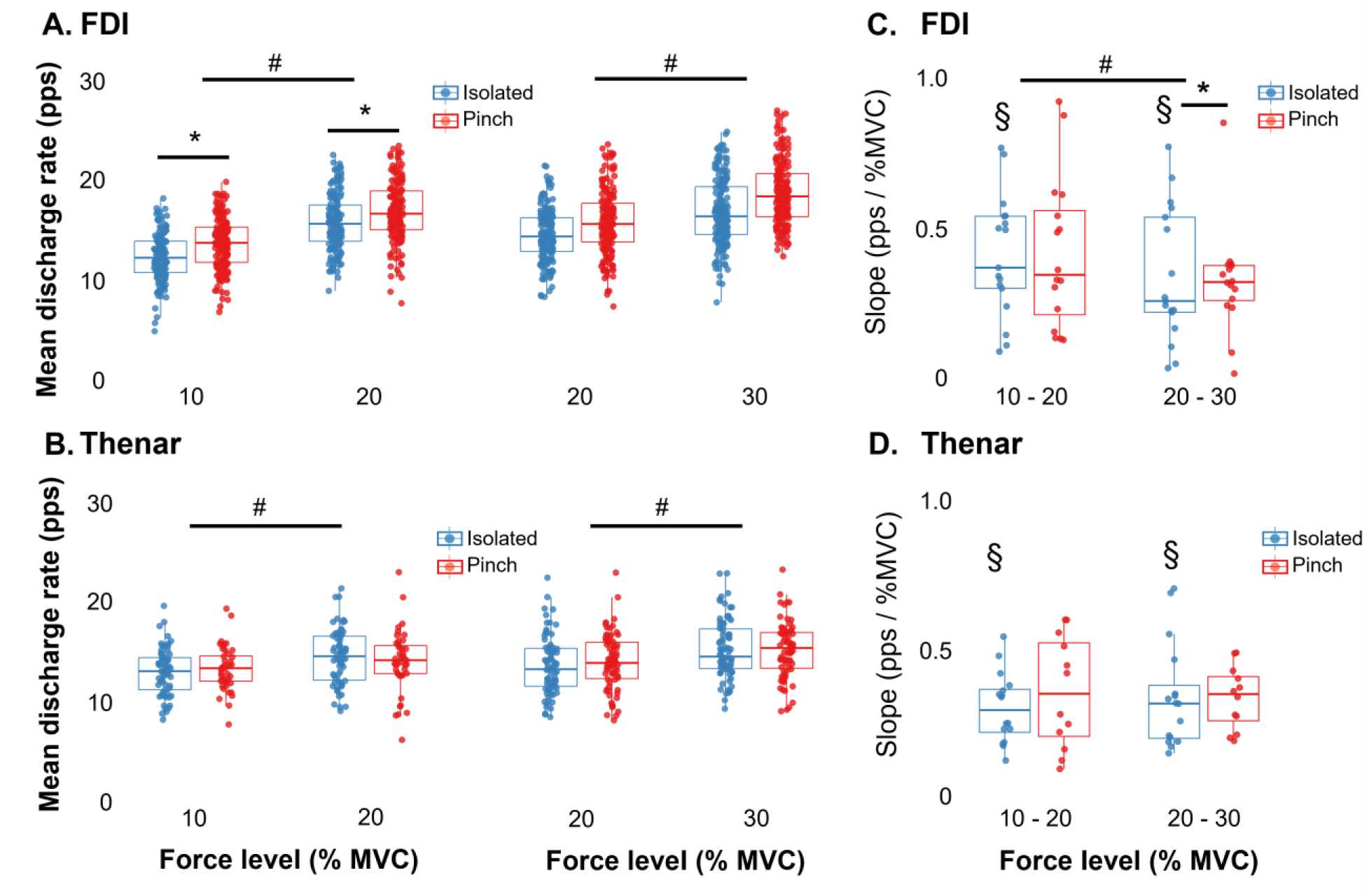
Mean discharge rate group results during plateau phase of isolated (blue) and pinch (red) tasks for the first dorsal interosseous (A) and thenar (B). In these results, motor units were tracked separately for 10-20% MVC and 20-30% MVC. Slope of linear regression across forces for the first dorsal interosseous (C) and thenar (D). Circles identify individual motor units. Horizontal traces, boxes, and whiskers denote the median value, interquartile interval, and distribution range. * Signifies significant difference between task p<0.05; # Signifies significant difference across force levels p < 0.05; § Signifies significant differences between muscle p<0.05.

A task-dependent effect was observed in the FDI. For motor units matched between 10-20% MVC, the mean discharge rate was higher during the pinch task than during isolated index finger flexion (F_(1,608.56)_ = 8.111, p = 0.005, **Figure 3A**). In addition, there was a significant muscle × task interaction of the slope of the mean discharge rate (10-20% MVC: F_(1,.367.67)_ = 7.366, p = 0.007; 20-30% MVC: F_(1,499.03)_ = 5.104, p = 0.024). Post hoc comparisons revealed that for the 20-30% MVC, the slope of the FDI was higher in the isolated finger flexion than in the pinch task (p < 0.001, **Figure 3C**). Across both force ranges, the slope of the FDI was also significantly higher than the slope of the thenar (10-20% MVC: F_(1,368.96)_ = 45.016, p < 0.001; 20-30% MVC: F_(1,454.60)_ = 46.787, p < 0.001, **Figure 3C and 3D**).

During the recruitment phase, the thenar exhibited higher mean discharge rates in the pinch task compared to isolated thumb flexion for the motor units matched between 20-30% MVC (F_(1,290.73)_ = 4.145, p = 0.043, see supplementary file). In contrast, no significant differences in mean discharge rate were observed during the de-recruitment phase under any condition (see supplementary file).

#### Matched units across 10-20-30% MVC

After completing the second step of tracking, identical motor units were identified across all three force levels. In the FDI, mean discharge rate significantly increased with force during both the recruitment phase (F_(1,532.10)_ = 13.412, p < 0.001, **Figure 4A**) and the plateau phase (F_(1, 728.04)_ = 306.040, p < 0.001, **Figure 4B**). In contrast, the thenar showed a significant increase only during the plateau phase, from 10% to 30% MVC (p < 0.001, **Figure 4B**).

**Figure 4.**
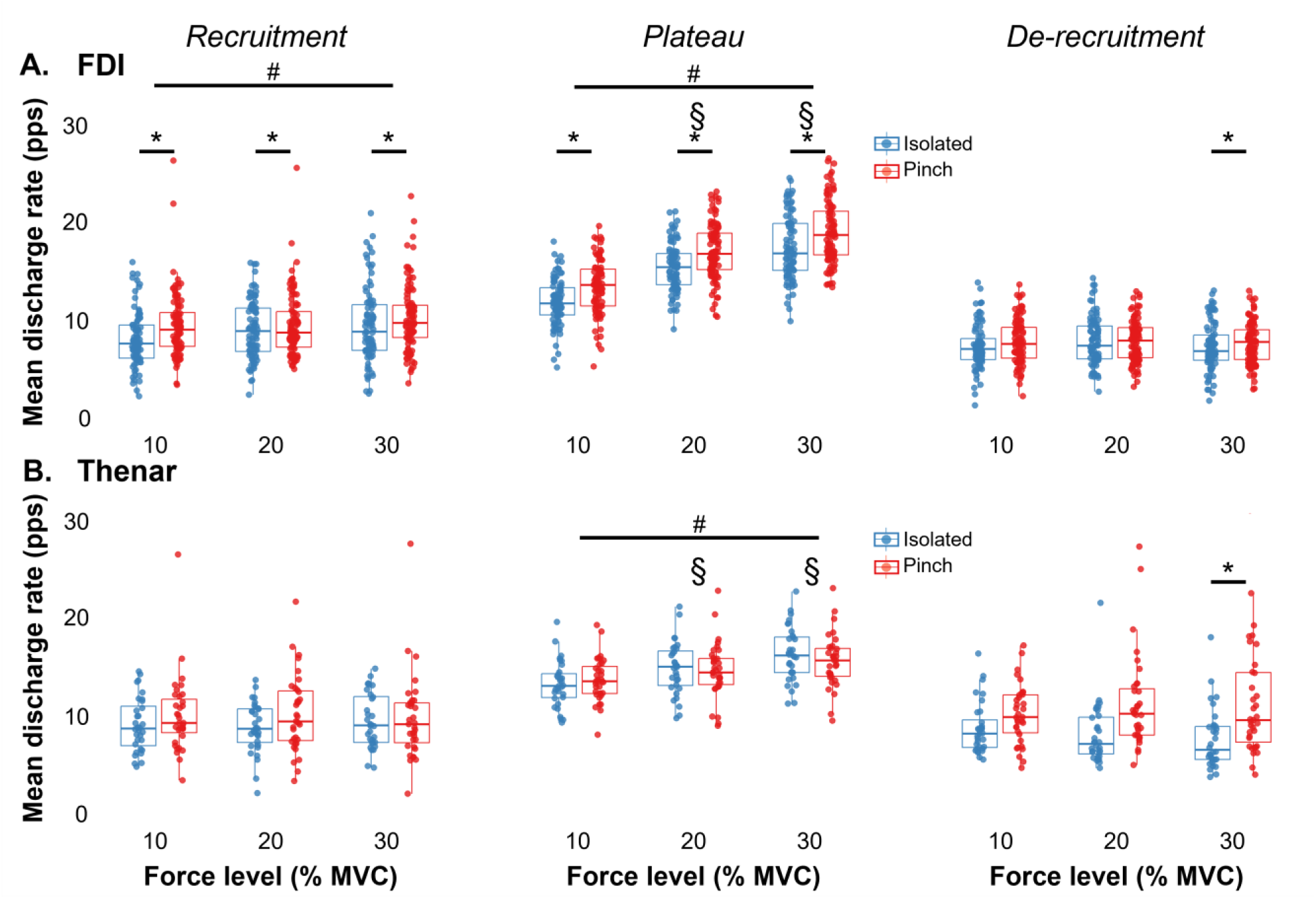
Mean discharge rate group results during recruitment (left), plateau (middle), and de-recruitment (right) phases of the isolated (blue) and pinch (red) tasks for (A) the first dorsal interosseous and (B) the thenar. In these results, motor units were tracked across 10-20-30% MVC. Circles identify individual motor units. Horizontal traces, boxes, and whiskers denote the median value, interquartile interval, and distribution range. * Signifies significant difference between task p<0.05; # Signifies significant difference across force levels p < 0.05; § Signifies significant differences between muscle p<0.05.

Task-dependent modulation of the motor unit discharge rate was also observed in the FDI. During the recruitment phase, mean discharge rate was higher in the pinch task than isolated finger flexion (F_(1,534.55)_ = 5.751, p = 0.017, **Figure 4A**). The same pattern was found during the plateau phase (F_(1, 532.96)_ = 9.546, p = 0.002, **Figure 4A**). During the de-recruitment phase, at 30% MVC, the FDI also had a higher discharge rate in the pinch task than the isolated contraction (p = 0.012, **Figure 4**).

Finally, a significant muscle × force interaction was observed during the plateau phase (F_(1, 728.04)_ = 45.531, p < 0.001). Post-hoc analysis revealed that at both 20% and 30% MVC, the FDI exhibited higher discharge rates than the thenar (p < 0.001).

### Motor unit recruitment and de-recruitment thresholds

#### Matched units between 10-20% and 20-30% MVC

For motor units matched between 10-20% and 20-30% MVC, recruitment threshold increased significantly with force level in both the FDI (10-20% MVC: p < 0.001; 20-30% MVC: p= 0.013, **Figure 5A**) and the thenar (10-20% MVC: p = 0.011; 20-30% MVC: p = 0.005, **Figure 5C**).

**Figure 5.**
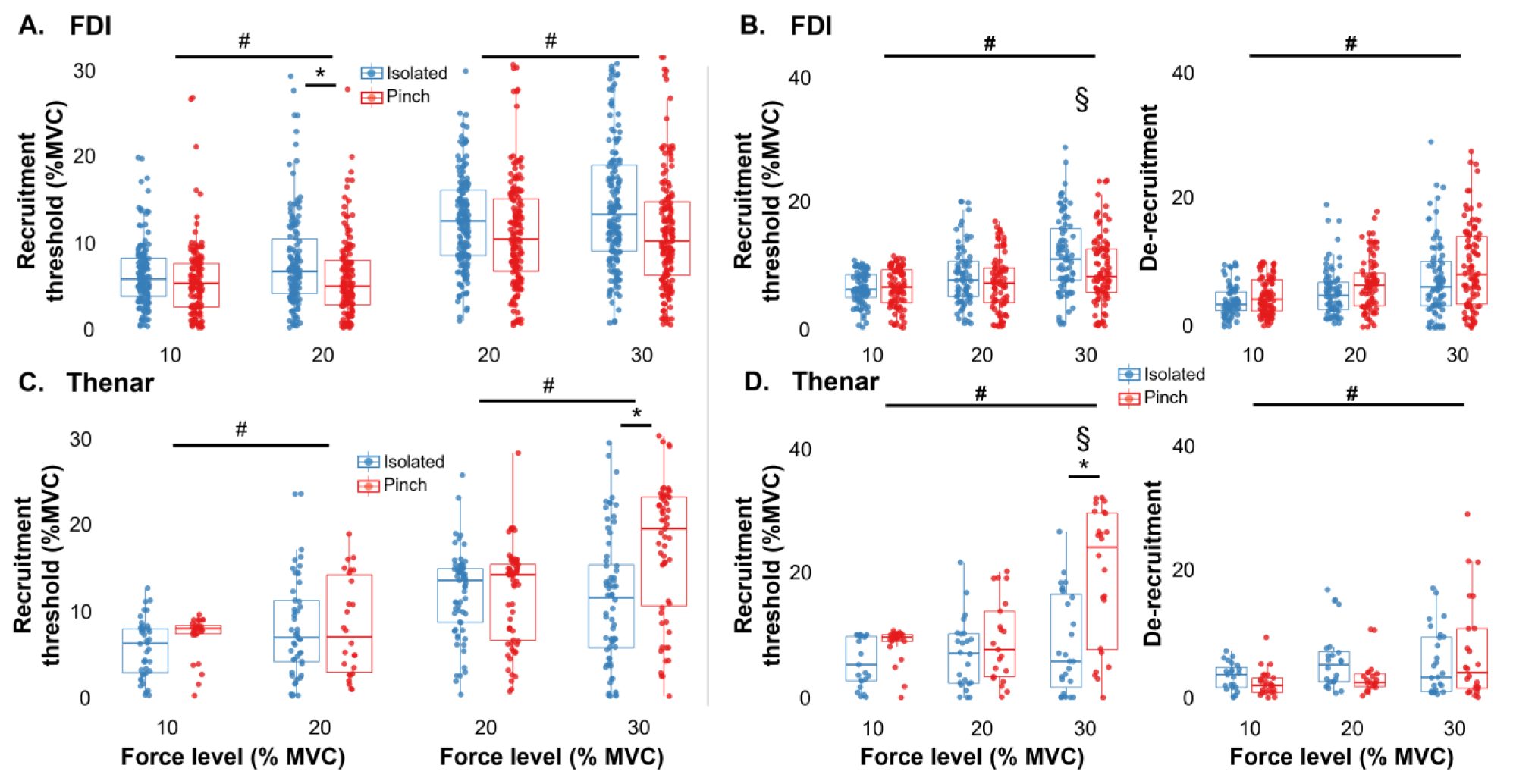
Motor unit recruitment threshold group results for the first dorsal interosseus (top) and the thenar (bottom) muscles. Results are displayed for motor units tracked separately for 10-20% MVC and 20-30% MVC (left) and across 10-20-30% MVC (right). Circles identify individual motor units. Horizontal traces, boxes, and whiskers denote the median value, interquartile interval, and distribution range. * Signifies significant difference between task p<0.05; # Signifies significant difference across force levels p < 0.05; § Signifies significant differences between muscle p<0.05.

In the thenar, 20-30% MVC matched units showed a significant main effect of task (F_(1,246.78)_ = 8.537, p = 0.004) and a force × task interaction (F_(1,244.87)_ = 13.617, p < 0.001). Post hoc analysis revealed higher recruitment thresholds in the pinch task than isolated thumb flexion at 30% MVC (p < 0.001, **Figure 5C**). In contrast, for 10-20% MVC matched units in the FDI, a task × force interaction was observed (F_(1,557.52)_ = 6.159, p = 0.013, **Figure 5A**), with post hoc tests indicating increased recruitment thresholds during isolated finger flexion than during the pinch task at 20% MVC (p < 0.001, **Figure 5A**). De-recruitment thresholds also increased with force level in both the FDI (10-20% MVC: p < 0.001; 20-30% MVC: p = 0.058) and thenar (10-20% MVC: p = 0.01; 20-30% MVC: p = 0.023). In the thenar, task (F_(1,241.663)_ = 5.193, p = 0.024) and task × force interaction (F_(1,240.36)_ = 5.384, p = 0.021) effects were observed for 20-30% MVC matched units, but post hoc comparisons did not reveal significant differences (see supplementary file).

#### Matched units across 10-20-30% MVC

For motor units tracked across all three force levels, recruitment threshold increased progressively with ascending force (F_(1,643.23)_ = 80.751, p < 0.001, **Figure 5B** and **5D**). Significant interactions were observed for force × task (F_(1,639.20)_ = 5.287, p = 0.022), force × muscle (F_(1,642.49)_ = 4.952, p = 0.026), and force × task × muscle (F_(1,639.32)_ = 18.126, p < 0.001). Post hoc comparisons revealed that at 30% MVC, the thenar had higher recruitment thresholds than the FDI (p < 0.001), and within the thenar, recruitment thresholds were higher in the pinch task than isolated conditions (p < 0.001, **Figure 5D**). For the de-recruitment threshold, only a main effect of force was observed, with values increasing with force (F_(1,638.91)_ = 52.451, p < 0.001, **Figure 5B** and **5D**).

## Discussion

This study investigated the motor unit discharge behaviour of the FDI and thenar during isometric contractions by tracking the same motor units across force levels (10%, 20%, and 30% MVC) during both an isolated and a tip pinch task. Three main findings were observed. First, the mean discharge rate increased with force in both muscles, but the slope of increase was steeper in the FDI, particularly between 10% and 20% MVC. Second, the FDI exhibited a lower recruitment threshold than the thenar, yet consistently higher mean discharge rates during the plateau phase. Third, motor unit behaviour was task-dependent. In the FDI, discharge rates were higher during the pinch than during isolated contractions, whereas in the thenar, task differences were limited and mainly observed at higher forces. Together, these results demonstrated that both muscle-specific properties and task demands influence force gradation in intrinsic hand muscles, highlighting distinct neural strategies for the FDI and thenar when acting independently or synergistically.

Motor unit behaviour in both the FDI and thenar followed a force-dependent pattern during submaximal contractions (from 10 to 30% MVC). Mean discharge rate and recruitment threshold increased progressively with force in both the isolated (finger flexion and thumb flexion) and pinch tasks. These results are consistent with the principle that greater force requirements are met through both the activation of additional motor units and modulation of their discharge rates (Fuglevand et al., 1993; Inglis et al., 2025; Maffiuletti et al., 2016; Milner-Brown et al., 1973). Interestingly, as force levels increase during isolated muscle contraction, the discharge rate does not rise in a fully linear manner. In the index finger contraction, the slope of the increase in FDI discharge rate was steeper between 10-20% MVC than between 20-30% MVC, whereas in the thenar of thumb flexion, the slopes did not differ significantly across force ranges. This trend suggests that the FDI relies more on rate modulation at lower force levels, which aligns with De Luca & Hostage (2010), who reported a similar slope of the FDI discharge rate across several force levels, while the thenar maintains a more uniform increase in discharge behaviour across force levels. These differences indicate that force-dependent increase in mean discharge rate reflects a general neural strategy to cope with increasing force demands, yet are fine-tuned differently across muscles.

Despite shared force-dependent adaptations, the FDI and thenar motor units displayed distinct discharge behaviours. The FDI consistently discharged at higher rates than the thenar during the plateau phase, with steeper discharge rate slopes across force levels. This observation aligns with the shallower ‘firing rate-force’ regression slope in FDI, with force increasing (Hu et al., 2014). Since most of the FDI motor units are recruited below 30% MVC, the elevated discharge rate of the FDI reflects greater rate modulation of its already-recruited motor units (De Luca et al., 1996; Kukulka & Clamann, 1981). Notably, the thenar is a muscle group rather than a single muscle, comprising the abductor pollicis brevis, flexor pollicis brevis, and opponens pollicis (Gupta & Michelsen-Jost, 2012). Functionally, the flexor pollicis brevis and opponens pollicis contribute more to pinch/grasp force, while the abductor pollicis brevis acts as a stabilizer engaged mainly for additional strength (Cooney et al., 1985). This might suggest that force generation in the thenar relies more on recruiting motor units across its subcomponents than increasing the discharge rate of individual units, which may explain its lower mean discharge rate compared to the FDI. Alternatively, this difference may reflect evolutionary adaptation, as the FDI is specialized for fine motor control (e.g., precise finger movements), where higher discharge rates enable more rapid and incremental adjustments in force (De Luca et al., 1982; Enoka & Fuglevand, 2001).

Task-specific differences further highlight these divergent strategies between the FDI and thenar. In the pinch task, the FDI was discharged at higher rates during both recruitment and the plateau phases. Interestingly, during de-recruitment the trend reversed, with the thenar exhibiting higher discharge rates than the FDI. This indicates that the two muscles adopt different rate coding strategies within the same synergistic task to meet dynamic demands. It could be explained by greater independence in the intrinsic muscle, which has also been found to exhibit less synchrony between adductor pollicis and FDI during the precision grip task (McIsaac & Fuglevand, 2008). The discharge behaviour of the thenar aligns with previous findings that the abductor pollicis brevis achieves optimal force at approximately 15-20 Hz during steady contractions (Howells et al., 2006), reflecting consistent firing rather than maximal rate modulation. These findings suggest that while both muscles adapt to force demands, the FDI emphasizes rapid rate modulation consistent with its role in fine motor control. However, the thenar relies more on recruitment across its subcomponents and stable firing, supporting its role in thumb stabilization and coordination during a pinch task.

The pinch task revealed further divergence in control strategies of the FDI and thenar relative to the isolated thumb and finger contractions. In the FDI, discharge rates were consistently higher during the pinch task than during isolated contractions across all phases (recruitment, plateau, and de-recruitment). Recruitment thresholds also differed by task: at 20% MVC, thresholds were higher during the isolated contractions than during the pinch task. This behaviour suggests that pinch tasks place greater demands on the FDI’s low-threshold motor units, leading to both higher discharge rates and lower recruitment thresholds relative to isolated contractions. In contrast to the FDI, the thenar exhibited task-dependent changes only during recruitment at 20-30% MVC, with discharge rates greater during the pinch task than during the isolated contraction. The thenar also had higher recruitment thresholds in the pinch task at 30% MVC, and across all pinch task conditions, the thenars thresholds were consistently higher than those of the FDI. Because higher thresholds require greater excitability to activate motor units, it implies that the thenar may be recruited later than the FDI during the pinch task. Notably, both the FDI and thenar muscles operated under identical force requirements in this study, ruling out “force level” as a confounding factor and reinforcing task-specific neural strategies. Anatomically, the FDI is a bipennate muscle located between the first and second metacarpal bones, functioning to abduct the index finger and assist the thumb during pinching movement (Bilbo & Stern, 1986; Infantolino & Challis, 2010). This makes the FDI a key contributor to both isolated index finger flexion and the pinch task. Indeed, previous EMG evidence supports that, during the pinch task, the adductor pollicis and FDI act as the primary force-generating muscles, while the opponens pollicis contributes as a secondary muscle whose activity increases as force demands rise (Cooney et al., 1985). This activation pattern suggests that the FDI acts as a ‘task initiator’ that drives early force generation during a pinch task, whereas the thenar contributes more to support activation and sustain increased force output.

The convergence of force-, muscle-, and task-dependent effects highlights that the FDI and thenar play complementary yet distinct roles in hand function. The FDI primarily adopts a “rate-based strategy”, rapidly increasing firing frequency to fine-tune force across both isolated and synergistic (pinch task) actions. Although the number of tracked motor units is limited in the thenar, this muscle seems to rely more on a “recruitment-based strategy”, particularly at higher force levels (30% MVC) or during a synergistic pinch task. Neural mechanisms likely reinforce this division through shared afferent input between the two muscles. For instance, electrical stimulation of the median nerve, which innervates the thenar, evokes a heteronymous H-reflex in the FDI, despite FDI not being directly innervated by the median nerve (Baudry et al., 2009). This finding confirms common afferent modulation across the two motor neuron pools, which may fine-tune their task-dependent responses. These observations align with broader motor control frameworks. Hug et al. (2023) emphasized that integrating Henneman’s size principle with common synaptic input reduces the computational load of controlling motor neuron pools. For the FDI and thenar, such a framework explains why their distinct strategies are not mutually exclusive but complementary, supporting both precision and stable force output during specific hand tasks.

A key limitation of this study is the relatively small number of tracked motor units obtained from the thenar across forces (10%, 20%, and 30% MVC). On average, only 3-4 matched motor units per participant were available for statistical analysis, which was substantially fewer than the tracked motor units in the FDI. Although this reduced number may have affected statistical power and representativeness of our findings, the decomposed motor unit yield in the thenar are in line with previous studies (Del Vecchio et al., 2023; Tanzarella et al., 2021).

Overall, the findings highlight distinct neuromuscular strategies employed by intrinsic hand muscles to meet increasing force demands. In conclusion, by tracking motor units in the FDI and thenar across three submaximal force levels (10%, 20% and 30% MVC), this study demonstrated that both mean discharge rate and recruitment threshold increased with force in isolated and pinch tasks. However, task- and muscle-specific differences emerged. The FDI, as the preliminary muscle in the pinch task, initiates the task, and its motor units’ behaviour relied on a rate-based strategy to modulate already active units. On the other hand, the thenar muscle involved several muscle groups that recruited additional units to achieve similar force outputs. These findings suggest that the FDI and thenar serve specific and complementary roles in hand motor control, optimizing the balance between precision and stability required for functional tasks such as pinch.

## Acknowledgements

This study was funded by the European Research Council Consolidator Grant INcEPTION contract no. 101045605. J Greig Inglis was supported by the Marie Skłodowska-Curie Actions Grant ‘MUDecomp’ agreement no. 101151712.

