## Supplementary file for "Motor unit rate coding in intrinsic hand muscles during isolated finger contractions and pinch task"

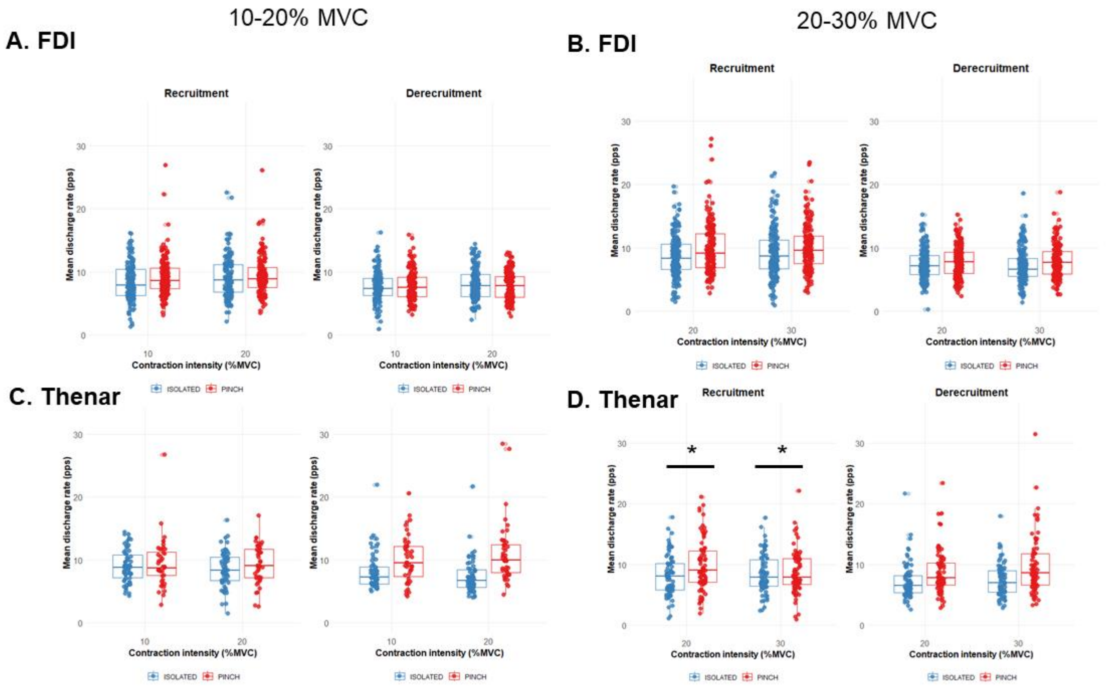


*Supplemental material 1: Mean discharge rate group results during recruitment and de-recruitment phases of isolated (blue) and pinch (red) tasks for the first dorsal interosseous (A)(B) and thenar (C)(D). In these results, motor units were tracked separately for 10-20% MVC and 20-30% MVC. Circles identify individual motor units. Horizontal traces, boxes, and whiskers denote the median value, interquartile interval, and distribution range. * Signifies significant difference between task $p<0.05$.*

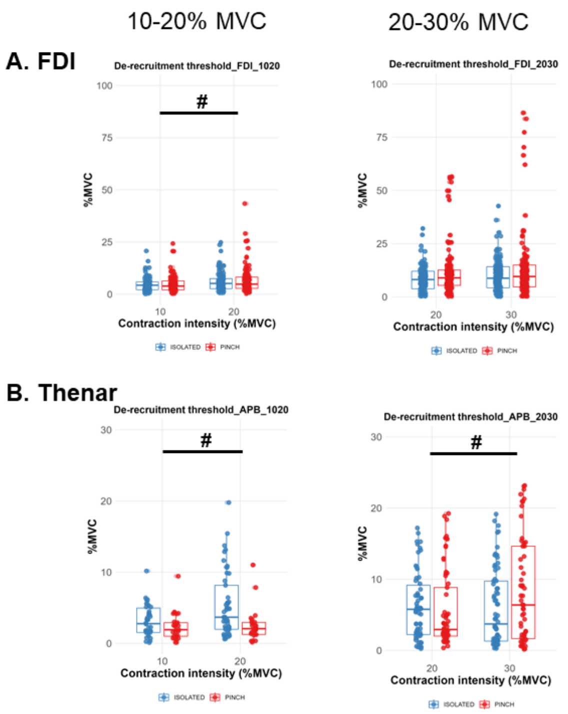


*Supplemental material 2: Motor unit de-recruitment threshold group results for the first dorsal interosseus (top) and the thenar (bottom) muscles. Results are displayed for motor units tracked separately for 10-20% MVC and 20-30% MVC (left) and across 10-20-30% MVC (right). Circles identify individual motor units. Horizontal traces, boxes, and whiskers denote the median value, interquartile interval, and distribution range. # Signifies significant difference across force levels $p < 0.05$.*
